# IgD^+^ Age-Associated B cells are the progenitors of the main T-independent B cell response to infection that generates protective Ab and can be induced by an inactivated vaccine in the aged

**DOI:** 10.1101/2022.02.04.479159

**Authors:** Olivia Kugler-Umana, Wenliang Zhang, Yi Kuang, Jialing Liang, Catherine H. Castonguay, Susan L. Tonkonogy, Ann Marshak-Rothstein, Priyadharshini Devarajan, Susan L. Swain

**Affiliations:** Department of Pathology, University of Massachusetts Medical School, Worcester, MA 01605, USA; College of Veterinary Medicine, North Carolina State University, Raleigh, NC 27607; Department of Medicine, University of Massachusetts Medical School, Worcester, MA 01605, USA

**Author notes:** **Corresponding author:** Susan L. Swain, Department of Pathology, University of Massachusetts Chan Medical School, Worcester, MA 01605, USA, Email ID, Priyadharshini Devarajan, Department of Pathology, University of Massachusetts Chan Medical School, Worcester, MA 01605, USA, Email ID.

**Keywords:** Age-Associated B cells, Aging, adaptive immunity, Influenza, TLR signals

## Abstract

Age-associated B cells (ABC) accumulate with age and are associated with autoimmunity and chronic infection. However, their contributions to acute infection in the aged and their developmental pathways are unclear. We find that the response against influenza A virus infection in aged mice is dominated by a Fas^+^GL7^-^ effector B cell population we call infection-induced-ABC (iABC). Most iABC express IgM and include antibody-secreting cells in the spleen, lung and bone marrow. We find that in response to influenza, IgD^+^CD21^-^CD23^-^ABC are the precursors of iABC. These IgD^+^ABC develop in germ free mice, so are independent of foreign antigen recognition, suggesting they arise from an intrinsic age-associated developmental program. The response of ABC to influenza infection, resulting in iABC, is T cell independent and requires both extrinsic TLR7 and TLR9 signals. In response to influenza infection, IgD^+^ABC can induce a faster recovery of weight and higher total anti-influenza IgG and IgM titers that can neutralize virus. Immunization with whole inactivated virus also generates iABC in aged mice. Thus, in unimmunized aged mice, whose other B and T cell responses have waned, IgD^+^ABC are likely the naïve B cells with the potential to become Ab-secreting cells and to provide protection from infection in the aged.

## Introduction

In elderly patients, lower respiratory tract infections caused by viruses such as influenza and SARS-CoV-2, are the most common cause of hospitalization (Ruuskanen et al. 2011; Kline & Bowdish 2016; Bartleson et al. 2021). In the US, over half of influenza-related hospitalizations and deaths and over 80% of COVID-related deaths, were in patients over the age of 65 (Kline & Bowdish 2016; Lees et al. 2020; Frasca & Blomberg 2020; Bartleson et al. 2021). Immune response to infections in aged populations are often weak and most current influenza vaccines fail to induce sufficient protective immunity to prevent severe illness (McElhaney et al. 2016; Bupp et al. 2018; Devarajan & Swain 2019; Henry et al. 2019; Frasca & Blomberg 2020). Defining the changes in the immune system with age and in response to influenza infection, is likely to provide insights into the heightened susceptibility of elderly patients to new infection with rapidly mutating viruses like influenza and other ssRNA viruses. Determining the mechanisms that promote the most effective immune responses in the aged, should inform strategies to improve vaccines and therapies for the aged.

The immune system undergoes many changes with age that contribute to reduced responses to infections and to immunizations (Frasca & Blomberg 2020; Shaw et al. 2013; Swain et al. 2021). Responses mediated by innate cells are reduced (Wong et al. 2017; Bupp et al. 2018; Shaw et al. 2013; Pillai et al. 2016) with age. Multiple steps in the development of adaptive responses of T and B cells are also impaired (Frasca & Blomberg 2009; Haynes & Swain 2006). Aged mice have a much smaller pool of naïve CD4 T cells and thus have a more restricted TCR repertoire (Dorshkind & Swain 2009). The aged naive CD4 cells respond less vigorously to antigen and develop fewer CD4 T cell effectors (Chinn et al. 2012; Zhang et al. 2014; Shifrut et al. 2013; Haynes et al. 2004; Jones et al. 2010), including an impaired T follicular helper (TFH) response (McElhaney 2011; Eaton et al. 2004). The aged effector T cells develop little T cell memory (McElhaney et al. 2016; Zhou & McElhaney 2011). Like T cells, the generation of immature B cells in the bone marrow (BM) decreases with age (Ademokun et al. 2010; Scholz et al. 2013), so the naïve mature B cell pool consists of longer-lived, older follicular B cells (FOB) and marginal zone B cells (MZB) (Guerrettaz et al. 2008; Scholz et al. 2013; Gibson et al. 2009) that respond less vigorously to antigenic challenge than B cells from younger individuals (Frasca & Blomberg 2009; Frasca & Blomberg 2020). In responses to new or mutated pathogens, these impairments in conventional naïve B cell and TFH responses cause decreased germinal center B cell (GCB) responses (Haynes et al. 2003; Haynes et al. 2004; Brahmakshatriya et al. 2017; Jones et al. 2010) with fewer isotype-switched Ab-secreting cells (AbSC), fewer long-lived PC (LLPC) and fewer memory B cells (Bmem) (Lefebvre et al. 2016; McElhaney et al. 2016; Zhang et al. 2014; Eaton et al. 2008; Brahmakshatriya et al. 2017; Lazuardi et al. 2005). This leads to less effective clearance of virus and less durable protective memory (Frasca & Blomberg 2020). Since responses of naïve B cells to novel antigens presented by new variants of viruses and vaccines are poor in the aged (Scholz et al. 2013; Frasca & Blomberg 2009), there is an increased reliance on previously generated memory B cell responses (Devarajan & Swain 2019). Although there are fewer naïve FOB cells with age, the total number of B cells is similar in young and aged mice (Scholz et al. 2013), raising the possibility that additional subsets of B cells might exist in aged animals.

Michael Cancro’s group identified a novel subset of splenic B cells that accumulated with age in unimmunized mice (Hao et al. 2011) and were defined by lack of expression of CD21 and CD23 (Hao et al. 2011; Ratliff et al. 2013). B cells of that phenotype were subsequently identified in humans (Rubtsov et al. 2011; Nipper et al. 2018) and were also called age-associated B cells (ABC). Experiments using ABC from unimmunized mice, indicated that addition of TLR7 or TLR9 stimulation paired with anti-µ activation was needed to induce their proliferation *in vitro* (Hao et al. 2011). Pippa Marrack and colleagues simultaneously described a murine CD21^-^ CD11b^+^CD11c^+^ population which increased with age and also was called ABC (Rubtsov et al. 2011). They found these CD11c^+^ABC accumulated in unimmunized aged and young autoimmune-prone mice (Rubtsov et al. 2011; Rubtsov et al. 2013; Manni et al. 2018) and that they played a key role in autoimmunity. The Marrack ABC resembled extrafollicular (EF) B cells that develop in autoimmunity and chronic infections described by others based on shared expression of CD11c and T-bet (Rubtsova et al. 2017; Jenks et al. 2018). Thus, there is compelling evidence that ABC are a unique B cell population that develops with age and that ABC play key roles in autoimmunity and are associated with aged responses to chronic infection in both mice and humans. However, the ABC characterized in chronically infected/autoimmune human patients most likely develop after exposure to foreign antigen, suggesting they differ from the ABC that develop in unimmunized aged mice that are expected to include naïve B cells. Therefore, cells called ABC may encompass B cells at multiple stages of differentiation (Kugler-Umana et al. 2020).

Indeed, Cancro has suggested that CD21^-^CD23^-^ABC are an antigen (Ag)-experienced memory B cell population (Cancro 2020; Russell Knode et al. 2017), since they require signals from CD40L and thus likely cognate interaction with CD4 helper T cells for their development (Russell Knode et al. 2017) and collectively contain more somatic mutations in their B cell receptors than naïve follicular B cells (Russell Knode et al. 2017). However, we note that in unimmunized aged mice the majority of ABC still express surface IgD (Hao et al. 2011; Swain et al. 2017) and express a diverse B cell receptor repertoire (Russell Knode et al. 2017), a phenotype more compatible with a naïve population of B cells (Hao et al. 2011).

The extent of the role of ABC in acute infections and their contribution to anti-pathogen immunity are not well-defined. Given that the ABC population becomes a larger B cell subset as mice get older, making up more than one third of B cells in the aged mice (18-20 mo) (Hao et al. 2011), and that FOB at this age respond very poorly (Frasca & Blomberg 2020; Scholz et al. 2013), it seems likely that in the aged, ABC make an important contribution to protective immunity and provide an alternate pathway to Ab production to pathogens. Thus, we set out to define the subset of ABC in unimmunized aged mice that responds to infection, the mechanisms that drive their response and whether they give rise to protective AbSC that combat infection.

We analyzed the kinetics of B cell responses following influenza A virus (IAV) infection of aged vs young mice (Brahmakshatriya et al. 2017; Lefebvre et al. 2016) and found that aged mice developed an effector B cell population that expressed Fas but not GL7 (iABC). This population was the dominant B cell response in aged mice with a response that peaked at 21 days post infection (dpi) in the spleen, BM, and lung. Many of the iABC became AbSC (CD138^+^) that expressed IgM, with small subsets expressing IgG2b and IgA. The CD21^-^CD23^-^ABC in aged naïve wild type mice included both surface IgD^+^ and IgD^-^ B cells, and we found the IgD^+^ABC are the precursors of the great majority of iABC generated during influenza infection. The generation of iABC from IgD^+^ABC did not require T cell help but required both extrinsic TLR7 and TLR9 signals. Unlike iABC, the development of precursor ABC in aged mice did not require endosomal TLR and occurred in germ-free (GF) mice in the absence of foreign Ag. This suggests that ABC development is part of an intrinsic program of aging and that IgD^+^ABC are most likely naive cells. We found that IgD^+^ABC-derived AbSC can make influenza-specific IgM and IgG Ab, which can neutralize influenza and can accelerate recovery from infection-induced weight loss. Importantly, immunization with whole inactivated PR8 influenza virus (WIV), generated an equivalent iABC response in aged mice. Thus, we speculate that in response to infection, the IgD^+^ABC population in aged individuals generates AbSC that provide Ab-mediated protection and that in the aged ABC provide an alternate B cell pathway to protect against novel influenzas, SARS-CoV2 and other viruses that depend on naïve B cell responses.

## Results

### In aged mice, iABC are the dominant responding B cells to IAV

To better understand the features that distinguish B cell response in young (3-4 months) and aged (18-20 months) C57BL/6 mice, we monitored the *in vivo* kinetics of their B cell response to IAV infection. The young mice developed a robust GCB population, defined by expression of both Fas and GL7, that peaked at 14 dpi and was found in the spleen, but not lung or bone marrow (BM) (Fig.1a, S1a) (Lefebvre et al. 2016). In contrast, in aged mice we found very few GCB (Fig.1a, S1a) and instead they developed a dominant Fas^+^ GL7^-^ population, which we termed infection induced age-associated B cells (iABC). The iABC peaked at 21 dpi and were found in spleen, lung and BM (Fig.1a, S1a). As expected, neither iABC nor GCB developed in the absence of IAV infection (0 dpi) (Fig.1a). These results demonstrate that in naïve aged mice infected with IAV, generation of iABC is the dominant outcome of the B cell response and hence the most likely source of potentially protective Ab.

**Figure 1:**
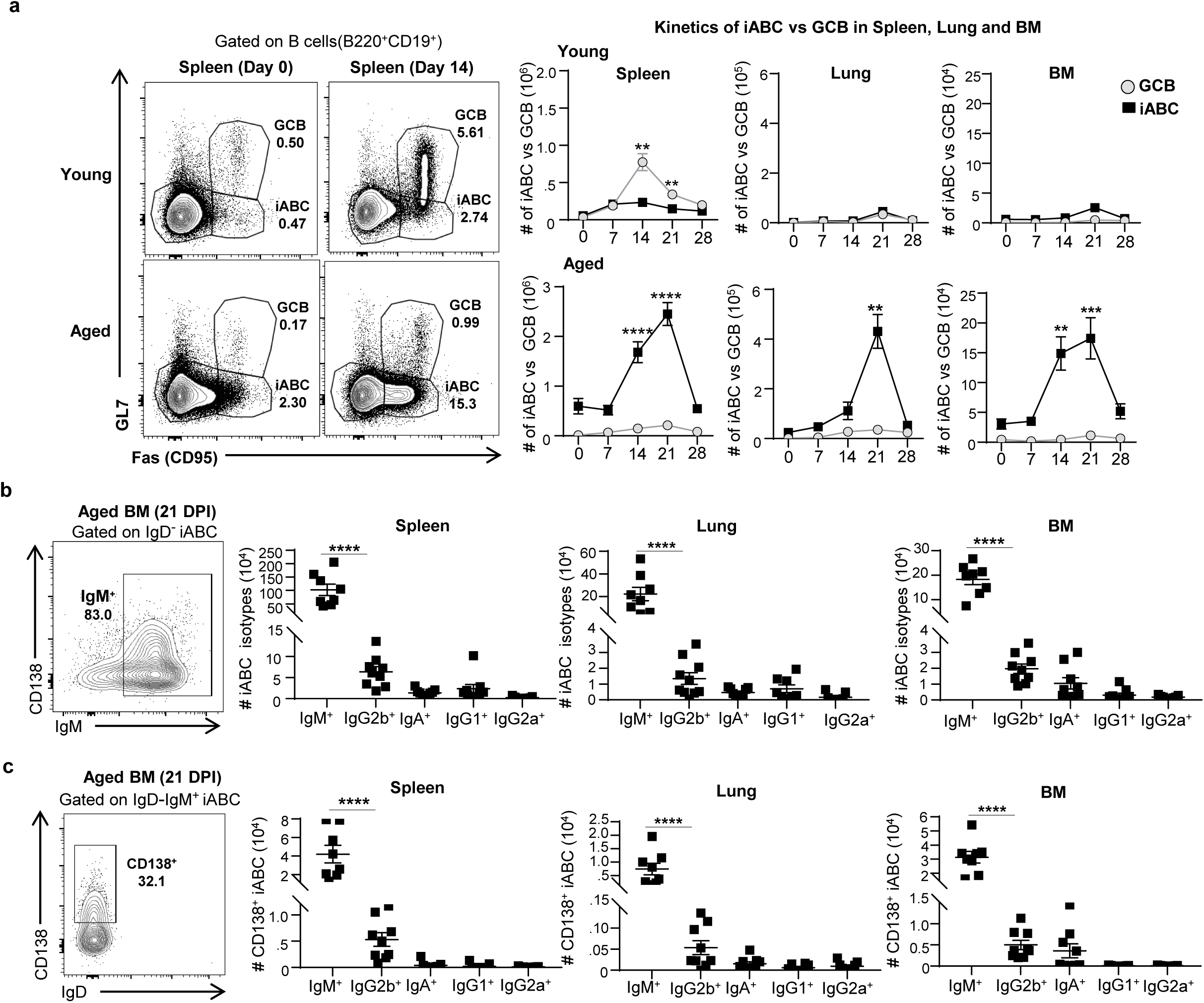
Kinetics of Aged B cell Effector and Ab Response. <u>Experimental Design:</u> Aged (18-20 M) and young (3-4 M) B6 mice, were infected i.n. with 0.3 LD50 PR8 (IAV) and sacrificed at 0, 7, 14, 21, and 28 dpi for analysis of B cell responses. Spleen, lungs and bone marrow (BM) were harvested at each time point and analyzed by flow cytometry. (a) Representative FACS plots indicate expression of Fas and GL7 by gated B cells (left) and the kinetics of iABC vs. GCB response is expressed as total cell numbers per organ of iABC (Fas^+^GL7^-^) and GCB (Fas^+^GL7^+^) in spleen, lung and BM of young and aged mice. (b) Representative FACS plots and total numbers of IgM^+^, IgG2b^+^, IgA^+^, IgG1^+^, and IgG2a^+^ iABC in spleen, lung and BM at 21 dpi. (c) Representative FACS plots and total numbers of IgM^+^CD138^+^, IgG2b^+^CD138^+^, IgA^+^CD138^+^, IgG1^+^CD138^+^, and IgG2a^+^CD138^+^ iABC in spleen, lung and BM at 21 dpi (n=6-12 pooled from 2-3 separate experiments) Error bars represent s.e.m. Statistical significance determined by two-tailed, unpaired Student’s t-test; \**p* < .05; \*\**p* < .01; \*\*\**p* < .001, \*\*\*\**p* < .0001

### iABC include CD138^+^ plasma cells, that express mostly IgM with smaller IgG2b^+^ and IgA^+^ subsets

We asked if iABC include isotype-switched B cells that express an AbSC phenotype (CD138^+^). We found that among responding iABC (IgD^-^, Fas^+^, GL7^-^) in the spleen, lung, and BM at 21 DPI, over 80% expressed intracellular IgM (Fig. 1b). About one-third of IgM^+^ iABC co-expressed CD138^+^ in the BM, indicating that IgM^+^ iABC include AbSC (Fig. 1c). Therefore, the largest fraction of the iABC generated by IAV infection are IgD^-^IgM^+^ responding B cells which have not switched to other isotypes. This is consistent with the fact that influenza-specific IgG antibodies decrease with age (Lefebvre et al. 2016). Further analysis of the iABC revealed a population of IgG2b^+^ iABC in the lung (11-15%) and spleen (2-7%), and smaller populations of iABC expressing IgG1(1-4%) and IgA (2-6%) in the lungs (Fig. 1b, Fig. S1b). The presence of AbSC among iABC in the lung is important since this is where influenza replicates. We did not detect any intracellular IgG2c (Fig. S1b). Amongst IgD^-^IgM^-^iABC in the BM of aged mice, we detected IgG2b^+^ iABC, as well as smaller populations of IgG1^+^ and IgA^+^ iABC (Fig. S1b). Thus, the iABC response to IAV infection can develop into AbSC of several isotypes in multiple sites including the lung and BM.

### ABC in unimmunized mice are a heterogenous population of IgD^+^ and IgD^-^ABC

Some studies have suggested that ABC are memory B cells that accumulate in aged mice (Russell Knode et al. 2017; Du et al. 2019; Cancro 2020). However our preliminary data suggested that the largest fraction of ABC express surface IgD (Hao et al. 2011; Swain et al. 2017), a phenotype more compatible with a naïve B cell population (Hao et al. 2011). To evaluate the heterogeneity of the total ABC population in unimmunized aged mice, we analyzed the surface IgD and IgM expression of CD21^-^CD23^-^ABC (Fig. 2a) and compared their expression with those of the two conventional naïve B cells present: FOB (CD21^+^CD23^+^) and MZB (CD21^+^CD23^-^) (Fig. 2b). A major population of ABC (about 66%) expressed surface IgD, as did almost all FOB (about 94%) and most MZB (about 84%) (Fig. 2b). This is consistent with the hypothesis that the population of ABC in unimmunized aged mice is composed mostly of naïve B cells. Unlike FOB and MZB, the ABC population also contained a substantial fraction of IgM^+^IgD^-^ B cells (about 21%) and IgM^-^IgD^-^ B cells (about 13%), indicating the ABC population is heterogenous and suggesting it might contain some B cells that are not naïve, some of which might be memory cells.

**Figure 2:**
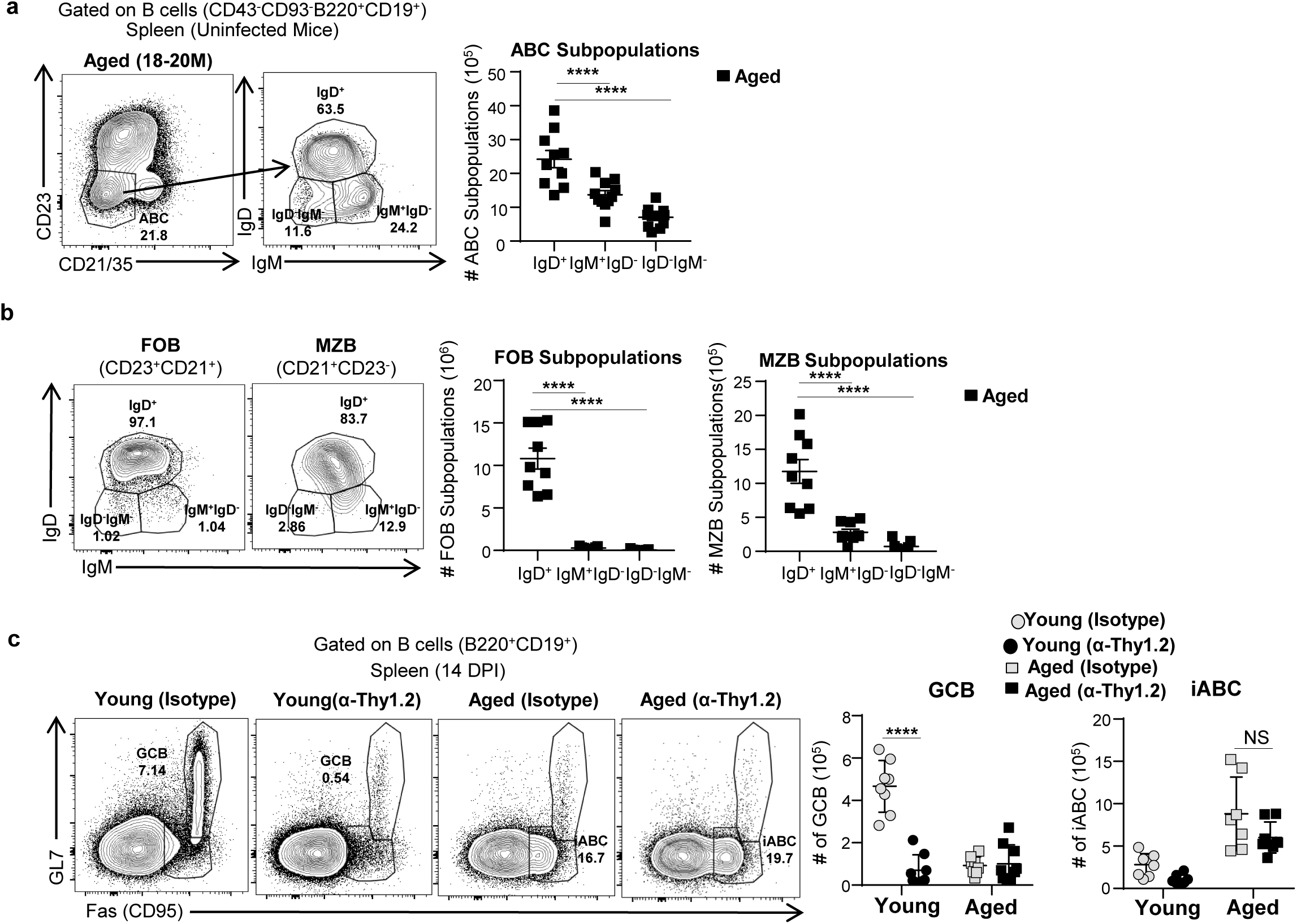
ABC are a heterogenous population and iABC develop independently of T cells-. <u>Experimental Design</u>: Uninfected Aged (18-20M) and young (3-4 M) female B6 mice were sacrificed. Spleens were harvested and CD23^-^CD21^-^ABC, FOB (CD23^+^CD21^+^) and MZB (CD21^+^CD23^-^) were analyzed by flow cytometry. (a) Representative FACS plot of gated ABC (CD43^-^CD93^-^B220^+^CD19^+^ CD23^-^CD21^-^) (left) and the number IgD^+^, IgM^+^IgD^-^ and IgD^-^IgM^-^ subsets in ABC, (b) Representative FACS plots and numbers of IgD^+^, IgM^+^IgD^-^, and IgD^-^IgM^-^ FOB (CD23^+^CD21^+^) and total IgD^+^, IgM^+^IgD^-^, and IgD^-^IgM^-^ MZB (CD21^+^CD23^-^) (right) in aged female B6 mice (n=9-12 pooled from 5-6 separate experiments) <u>Experimental Design:</u> Aged (18-20 M) and young (3-4 M) female B6 mice were treated with anti-Thy1.2 or isotype control Ab on 0 DPI and 7 DPI and infected with 0.3LD50 PR8 (IAV) and sacrificed at 14 dpi. Spleens were harvested and analyzed for iABC and GCB by flow cytometry as in Fig. 1. (c) iABC and GCB responses in aged and young mice depleted of T cells by Thy1.2 Ab. Representative FACS plots and cell numbers of iABC (Fas^+^GL7^-^) and GCB Fas^+^GL7^+^ gated on mature B cells (B220^+^CD19^+^) in aged and young mice in the spleen. (n=8-10 pooled from 2-3 separate experiments) Error bars represent s.e.m and statistical significance determined by two-tailed, unpaired Student’s t-test; \**p* < .05; \*\**p* < .01; \*\*\**p* < .001, \*\*\*\**p* < .0001

### iABC develop independently of T cells

We know aged mice have a substantially decreased CD4 T helper response (Lefebvre et al. 2016). Since aged mice nonetheless develop a robust iABC in response to influenza (Fig. 1), we hypothesized that this subset is less dependent on CD4 T cell help. To further evaluate this possibility, we depleted T cells using anti-Thy1.2 antibody (Strutt et al. 2012) and then infected young and aged mice with IAV. As expected, Thy1.2 depletion reduced the generation of GCB in young mice (Fig. 2c) compared to the controls (isotype control antibody-treated). In contrast, Thy1.2 depletion did not reduce iABC generation in aged mice (Fig. 2c). Therefore, iABC development in aged mice in response to IAV is T-independent.

### IgD^+^ ABC give rise to iABC

Since the CD21^-^CD23^-^ABC from aged mice are composed of IgD^+^ and IgD^-^ABC we asked which subpopulation contains the progenitors that give rise to iABC in response to influenza infection. We hypothesized that IgD^+^ABC would give rise to more iABC because they may be a naïve population, with a broad repertoire that would contain a higher fraction of B cells that could respond to IAV. We developed an adoptive transfer system using SAP^-/-^ hosts that are deficient in T cell help. SAP^-/-^ mice do not generate TFH responses (Kamperschroer et al. 2008; Kamperschroer et al. 2006), and thus mimic the aging host environment where few TFH develop in response to influenza infection (Lefebvre et al. 2016). We sorted CD45.2^+^, aged IgD^+^ABC and IgD^-^ABC (Fig. S2a) and transferred them into young allotype-marked (CD45.1) SAP^-/-^ hosts and infected with IAV (Fig. 3a). Since ABC have been implicated in autoimmunity (Phalke & Marrack 2018), we also transferred each subset to uninfected (no IAV) SAP ^-/-^ hosts to determine if the responses we generate were dependent on IAV infection (Fig. 3a). Analysis of donor populations, identified by CD45.2, at 21 dpi, revealed that donor IgD^+^ABC gave rise to an iABC population (Fas^+^ GL7^-^) in both the spleen and BM. In contrast, the IgD^-^ABC subset gave rise to only a very small iABC response (Fig. 3b,c). As expected, iABC were generated in IAV-infected but not uninfected mice (Fig. 3 b,c). There was a substantial population of donor cells that retained the naïve phenotype, expressing neither GL7 nor Fas (Fig. 3b) even in hosts that were infected, which is to be expected since we transferred a population of polyclonal naive ABC, of which only a very small fraction are expected to have a BCR that recognizes IAV.

**Figure 3:**
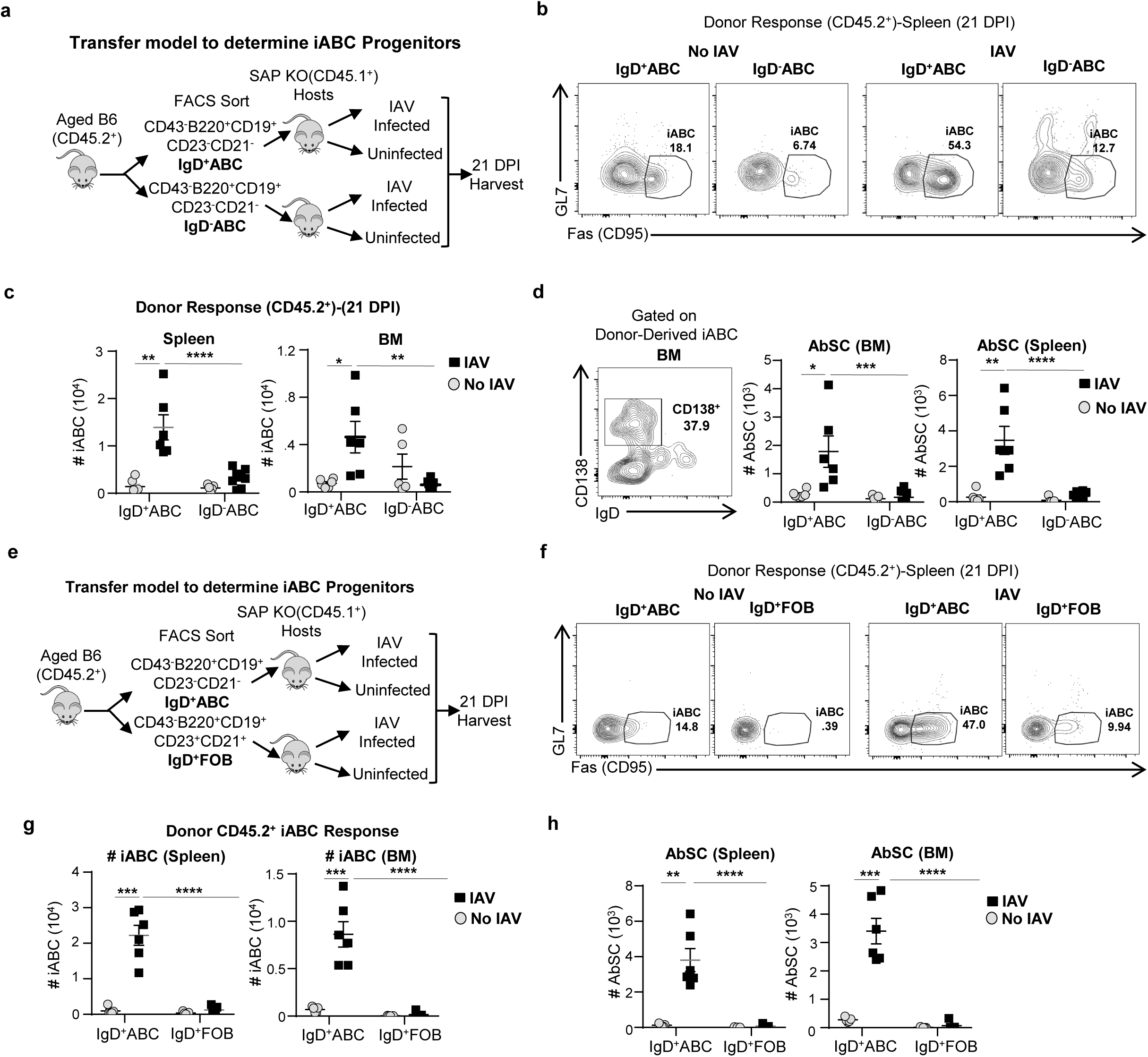
IgD^+^ ABC give rise to most of the iABC compared to IgD^-^ABC and FOB. (a) Schematic of the transfer model (left): Cells were enriched by positive MACS selection of cells stained with CD43 and Ter-119 and CD23 to remove immature B cells and FOB and B220^+^CD19^+^CD23^-^CD21^-^ mature B cells were sorted into IgD^+^ vs. IgD^-^ ABC by flow cytometry. These IgD^+^ and IgD^-^ ABC transferred into SAP^-^/^-^ CD45.1 hosts that were either infected with 0.4 LD50 PR8 (IAV) or left uninfected (No IAV). Hosts were sacrificed and spleens were harvested at 21 DPI (b) Representative FACS plots of iABC (Fas^+^GL7^-^) among gated donor B cells in hosts given IgD^+^ and IgD^-^ ABC. (c) Cell numbers of total donor iABC (CD45.2) derived from IgD^+^ vs. IgD^-^ ABC in the spleen and BM of infected vs. uninfected SAP^-^/^-^ hosts. (d) Representative FACS plots and cell numbers of total CD138^+^ iABC from IgD^+^ vs. IgD^-^ ABC in the spleen and BM of infected vs. uninfected SAP^-^/^-^ hosts. (e) Schematic of the transfer model: CD45.2 spleen cells were enriched by MACS Selection of CD43 andTer-119 cells and mature B cells (B220^+^CD19^+^) sorted into IgD^+^ABC (CD23^-^CD21^-^) and IgD^+^ FOB (CD23^+^CD21^+^) were sorted by flow cytometry and transferred into CD45.1 SAP^-^/^-^ hosts and either infected with 0.4 LD50 PR8 (IAV) or left uninfected (No IAV). Hosts were sacrificed and spleen and BM were harvested at 21 DPI. Representative FACS plots (f) and cell numbers (g) of total iABC from donor CD45.2 IgD^+^ABC, and IgD^+^FOB in the spleen and BM of infected vs. uninfected CD45.1 SAP^-^/^-^ hosts. (h) Cell numbers of total CD138^+^ iABC from IgD^+^ABC and IgD^+^FOB donors of infected vs. uninfected CD45.1 SAP^-^/^-^ hosts in the spleen and BM (n=5-6 pooled from 3-4 separate experiments). (n=5-6 pooled from 3-4 separate experiments) Error bars represent s.e.m Statistical significance determined by two-tailed, unpaired Student’s t-test; \**p* < .05; \*\**p* < .01; \*\*\**p* < .001, \*\*\*\**p* < .0001

To determine if the iABC derived from the transferred IgD^+^ABC donors become plasma cells, we stained for CD138 (Halliley et al. 2015; Jenks et al. 2018; Nutt et al. 2015) and analyzed the iABC that were AbSC by FACS. Following immunization, most of the iABC (80%) derived from the transferred IgD^+^ABC, had lost IgD expression, and a fraction expressed CD138^+^, indicating they were AbSC, in the spleen (28%) and BM (38%) (Fig. 3d). These data strongly suggest that in the aged IgD^+^ABC are the B cell precursors with highest potential to differentiate into the iABC and AbSC.

### IgD^+^ ABC, not FOB are the progenitors of iABC

Aged mice still maintain a substantial population of FOB and a small one of MZB cells (Hao et al. 2011), which could potentially give rise to iABC. Studies show that MZB responses are largely impaired due to several mechanisms in the aged (Birjandi et al. 2011; Turner & Mabbott 2017), making them unlikely to be precursors of iABC. However, if some FOB can respond in the aged, they might contribute to the AbSC response. We isolated equal numbers of IgD^+^ABC and IgD^+^ FOB, from aged, uninfected B6 mice, and transferred them into separate groups of infected or uninfected SAP^-/-^ CD45.1 hosts (Fig. 3e, S2c). After 21 dpi, we assessed the number of iABC formed from each of the donor subsets, in the spleen and bone marrow (BM) (Fig. 3f,g, S2d). Hosts that received IgD^+^ABC donor cells developed a substantial iABC response in spleen and BM, while FOB generated only background levels of iABC in the spleen and BM of the SAP^-/-^ hosts (Fig. 3f,g). None of the uninfected hosts developed significant iABC. Moreover, CD138^+^ iABC developed from IgD^+^ABC, but not from FOB in the spleen and in BM (Fig. 3h). These results strongly support the concept that in the aged mice, responses to IAV predominantly come from IgD^+^ABC that are the progenitors of an iABC response. The results indicate that in aged mice, the IgD^+^ABC are the progenitors of ABC that become iABC and AbSC in response to IAV infection.

### ABC accumulation is independent of commensal bacteria and foreign Ag

Little is known of the origins of the progenitor ABC that accumulate with aging. Previous studies suggested that Ag exposure plays a role in the generation of CD21^-^CD23^-^ABC (Russell Knode et al. 2017; Cancro 2020), but these studies did not separately evaluate the IgD^+^ progenitor subset and did not determine if Ag exposure was due to foreign or self Ag. We speculated that, like FOB progenitors, development of the IgD^+^ABC subset would not require foreign antigen stimulation, though we considered it might have a role in generation of the IgD^-^ subsets. To address the role of foreign and commensal Ag exposure, we compared ABC from our aged specific pathogen free (SPF) mice that contain Ag from commensal organisms and are exposed to foreign Ag from the environment, with aged germ-free (GF) mice that lack commensal microbes and are isolated throughout life to protect from exposure to environmental Ag. Aged female SPF and GF mice had equivalent high proportions and absolute numbers of splenic ABC (Fig. 4a). Like young SPF mice, young GF mice did not develop a distinct ABC population (Fig. 4a, S3a,b). We used surface staining to assess IgD^+^ and IgD^-^ ABC subsets within the ABC. The percentage and numbers of ABC that are IgD^+^, IgD^-^IgM^+^ and IgD^-^IgM^-^ were comparable in SPF and GF female mice (Fig. 4b). In both SPF and GF aged mice the IgD^+^ population was the largest population. We also compared the numbers of ABC in male and female mice. Aged male mice, both SPF and GF, had lower proportions of ABC compared to females, consistent with earlier indications that female mice develop more ABC (Hao et al. 2011; Russell Knode et al. 2017) (Fig. 4a). Thus, with age, accumulation of ABC occurs independent of exposure to foreign or commensal Ag and is most prevalent in female mice. This supports the concept that an intrinsic developmental program drives the development of ABC rather than any extrinsic factors such as environmental or commensal Ag stimulation.

**Figure 4:**
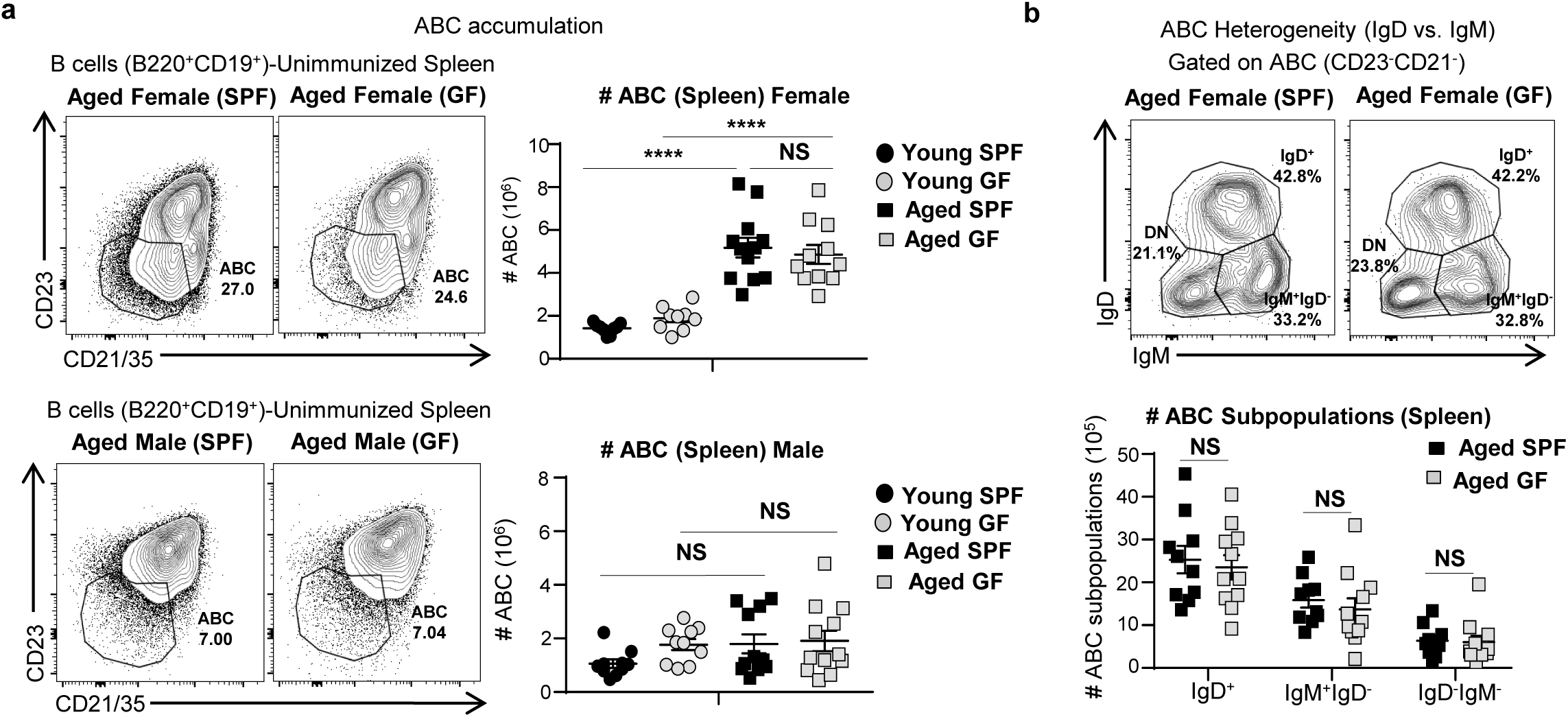
ABC subsets accumulate with age in germ-free (GF) as well as specific pathogen free (SPF) mice. <u>Experimental Design:</u> Unimmunized specific-pathogen free (SPF) and germ-free (GF), aged (18-20M) and young (3-4M) female and male B6 mice were sacrificed. Spleens were harvested and analyzed by flow cytometry. (a) Representative FACS plots and cell numbers of total ABC (CD21^-^ CD23^-^) in aged SPF vs GF B6 mice (b) Representative FACS plots and ABC containing IgD^+^, IgM^+^IgD^-^, and IgD^-^IgM^-^ subsets in aged SPF vs. GF female B6 mice (n=9-12 pooled from 7-8 separate experiments) Error bars represent s.e.m Statistical significance determined by two-tailed, unpaired Student’s t-test; \**p* < .05; \*\**p* < .01; \*\*\**p* < .001, \*\*\*\**p* < .0001

### Development of ABC is independent of TLR7 and TLR9

Previous studies have suggested that TLR7 plays a key role in the development of CD11c^+^ABC in autoimmune models and unimmunized aged mice (Rubtsov et al. 2013; Rubtsov et al. 2011; Naradikian, Hao, et al. 2016). To ask if progenitor ABC required TLR7 and TLR9 for their development, we examined the ABC population in aged TLR7^-/-^ and TLR9^-/-^ mice. ABC development was not impaired in either aged TLR7^-/-^ or aged TLR9^-/-^ mice compared to wild type (WT) aged mice (Fig. 5a, S3c). Thus, neither TLR7 nor TLR9 signaling is required for ABC accumulation with age. This further supports the concept that the progenitor ABC develop by an intrinsic program associated with aging, independent of external TLR7 and TLR9 stimuli.

**Figure 5:**
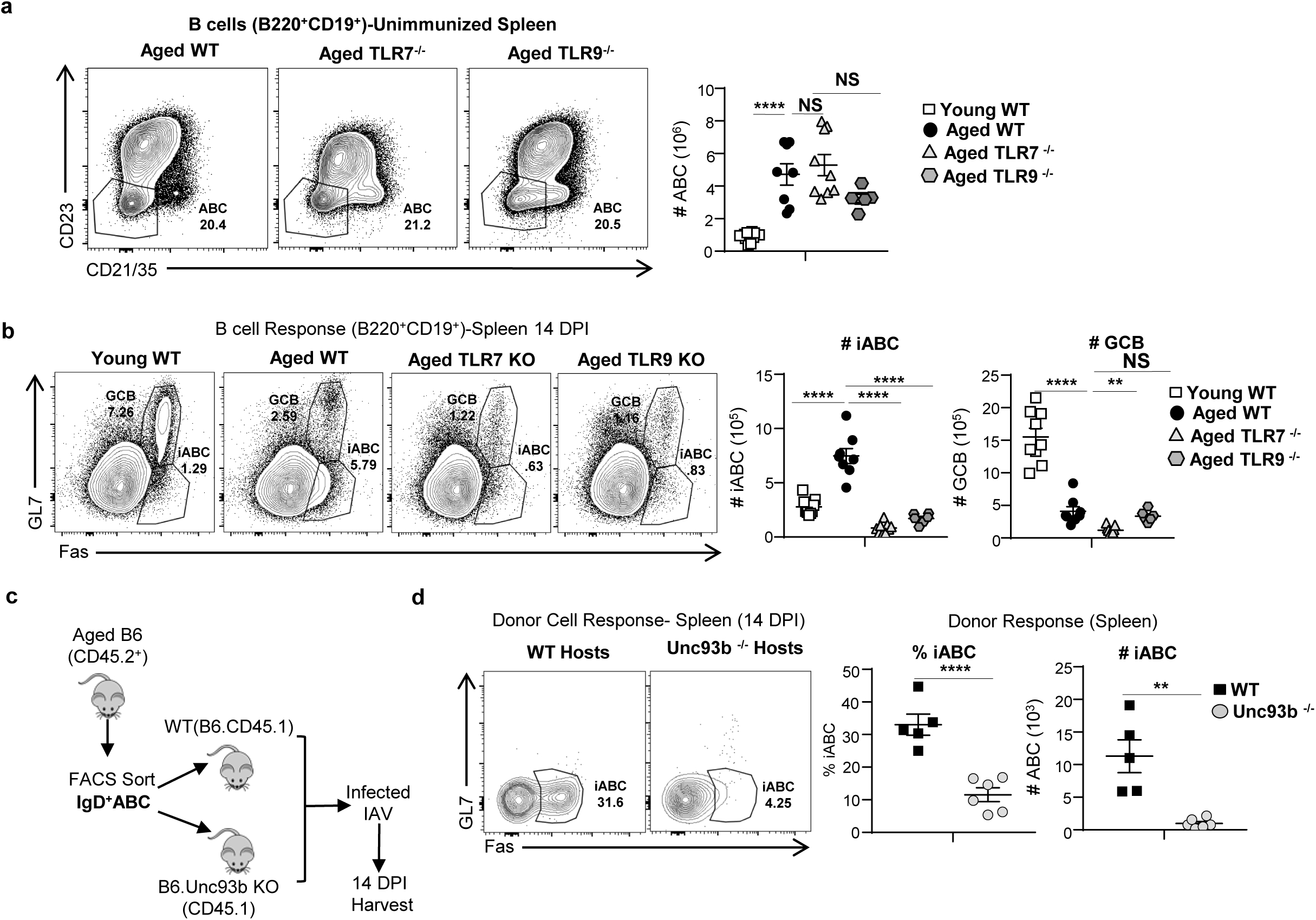
ABC develop independent of endosomal TLR signals, while iABC generation from ABC requires extrinsic endosomal TLR signaling. <u>Experimental Design</u>: Unimmunized Young, Aged WT, Aged TLR7 ^-^/^-^(*tlr*7^-/-^) and Aged TLR9 ^-^/^-^ (*tlr9* ^-^/^-^) female BALB/c mice were sacrificed. Spleens were harvested and B cells populations analyzed by flow cytometry. (a) Representative FACS plots and cell numbers of total ABC (CD21^-^ CD23^-^) in young and aged female BALB/c mice that were WT, aged TLR7 ^-^/^-^ and aged TLR9 ^-^/^-^ (n=5-11 pooled from 2-3 separate experiments). <u>Experimental Design:</u> Young and aged WT, aged TLR7 ^-^/^-^ and aged TLR9 ^-^/^-^ female BALB/c mice were infected with 0.4LD50 PR8 (IAV) and sacrificed at 14 dpi. Spleens were harvested and analyzed by flow cytometry. (b) Representative FACS plots and cell numbers of total iABC (Fas^+^GL7^-^) and GCB (Fas^+^GL7^+^) found in aged and young WT, aged TLR7 KO (*tlr*7^-/-^) and aged TLR9 KO (*tlr9* ^-^/^-^) female BALB/c mice (n=6-9 pooled from 2-3 separate experiments). (c) Schematic of the transfer model: IgD^+^ ABC (CD45.2) from aged B6 mice were isolated by sorting as in previous experiments and transferred into B6.CD45.1 WT or B6.CD45.1.Unc93b^-^/^-^ hosts, that were infected with 0.4 LD50 PR8 (IAV). Hosts were sacrificed and spleens were harvested at 14 dpi. (d) Representative FACS plots, percentage, and cell numbers of total iABC from donor CD45.2 IgD^+^ABC in the spleen of infected CD45.1 WT vs. Unc93b^-/-^ (*Unc93b*^-^/^-^) hosts. (n=5-6 pooled from 2-3 separate experiments) Error bars represent s.e.m Statistical significance determined by two-tailed, unpaired Student’s t-test; \**p* < .05; \*\**p* < .01; \*\*\**p* < .001, \*\*\*\**p* < .0001. NS=not significant.

### IgD^+^ABC require cell extrinsic endosomal TLR7 and TLR9 signals to generate iABC

Signaling through endosomal toll-like receptors (TLRs) and other pathogen recognition (PR) receptors has been shown to markedly enhance conventional anti-viral responses of FOB in young mice (Rawlings et al. 2012; Rookhuizen & DeFranco 2014; Iwasaki & Medzhitov 2015). *In vitro* TLR7 and TLR9 stimulation are needed to activate CD23^-^CD21^-^ABC isolated from unimmunized aged mice (Hao et al. 2011). To determine how much of a role TLR recognition pathways play in the generation of the iABC response to influenza infection, we infected TLR7 and TLR9 deficient aged mice and compared their responses to WT controls at 14 dpi. Neither TLR7^-/-^ nor TLR9^-/-^ developed iABC (Fig. 5b, S3d). The number of GCB was very low in all aged mice, as expected (Fig. 5b). Thus, unlike the progenitor ABC which develop independently of both foreign Ag and TLR7 and TLR9 pathways, the iABC response to influenza infection is highly dependent on both endosomal TLR7 and TLR9 pathways.

Most studies have focused on the cell-intrinsic role of TLR signals in ABC activation (Hao et al. 2011; Jenks et al. 2018; Cancro 2020; Rubtsov et al. 2013; Rubtsov et al. 2011; Teichmann et al. 2013). However, other studies have shown that TLR-dependent activation of dendritic cells can regulate autoreactive B cell response (Teichmann et al. 2013; Ols et al. 2016). Whether the ABC response depends only on direct TLR activation of the responding B cells (cell-intrinsic) or on TLR activation of non-ABC cells (cell-extrinsic) remains unclear. To evaluate a this, we transferred IgD^+^ABC from WT aged B6 mice into B6.Unc93b^-/-^ recipients (Fig. 5c). Unc93b is a chaperone protein required for the trafficking of TLR3, TLR7 and TLR9 receptors to the endocytic compartments where they engage their ligands, and thus Unc93-deficient mice cannot mount effective endosomal TLR responses (Pelka et al. 2018). The recipients were then challenged with IAV, and we analyzed iABC responses from donor cells at 14 dpi (Fig. 5d). Wild-type IgD^+^ABC donors did not develop iABC in Unc93^-/-^ hosts. This indicates that iABC generation from ABC progenitors requires extrinsic activation of TLR in host cells.

### ABC responses to IAV result in neutralizing anti-influenza Ab

In Fig.1, we showed that iABC are the major responding population to influenza infection in an aged immune response and that they become AbSC. To evaluate whether ABC-derived Ab can combat influenza infection in an adoptive transfer model, we needed a host with very limited B cell responses to influenza. The SAP^-/-^ mice used as hosts in previous figures (Fig. 3) still have B cells that can mount TFH-independent antibody responses (Crotty et al. 2003; Kamperschroer et al. 2006), so are not useful for assessing ABC-derived protection. We choose MD4 BCR transgenic mice whose B cells do not respond to IAV, because the MD4 receptor is specific for hen egg lysozyme (HEL) (Bortnick et al. 2012). MD4 hosts, not given ABC cells prior to infection, were used as negative controls (Fig. 6a). MD4 with transferred IgD^+^ABC, recovered more weight than control MD4 hosts without transferred cells (Fig. 6b). At 28-30 dpi, we collected serum to evaluate Ab levels and determine their ability to neutralize virus. MD4 hosts with ABC had higher titers of anti-influenza IgM and IgG in their serum compared to MD4 hosts without ABC (Fig. 6c,d). We used a plaque neutralization assay to determine if the serum Ab could neutralize influenza. The serum from MD4 hosts with transferred IgD^+^ABC neutralized IAV preventing plaque formation (Fig. 6e), while that from hosts without transferred ABC failed to do so. This suggests IAV-induced iABC make IAV-specific Ab responses, which can neutralize virus and hence have the potential to provide significant protection from influenza infection in aged animals.

**Figure 6:**
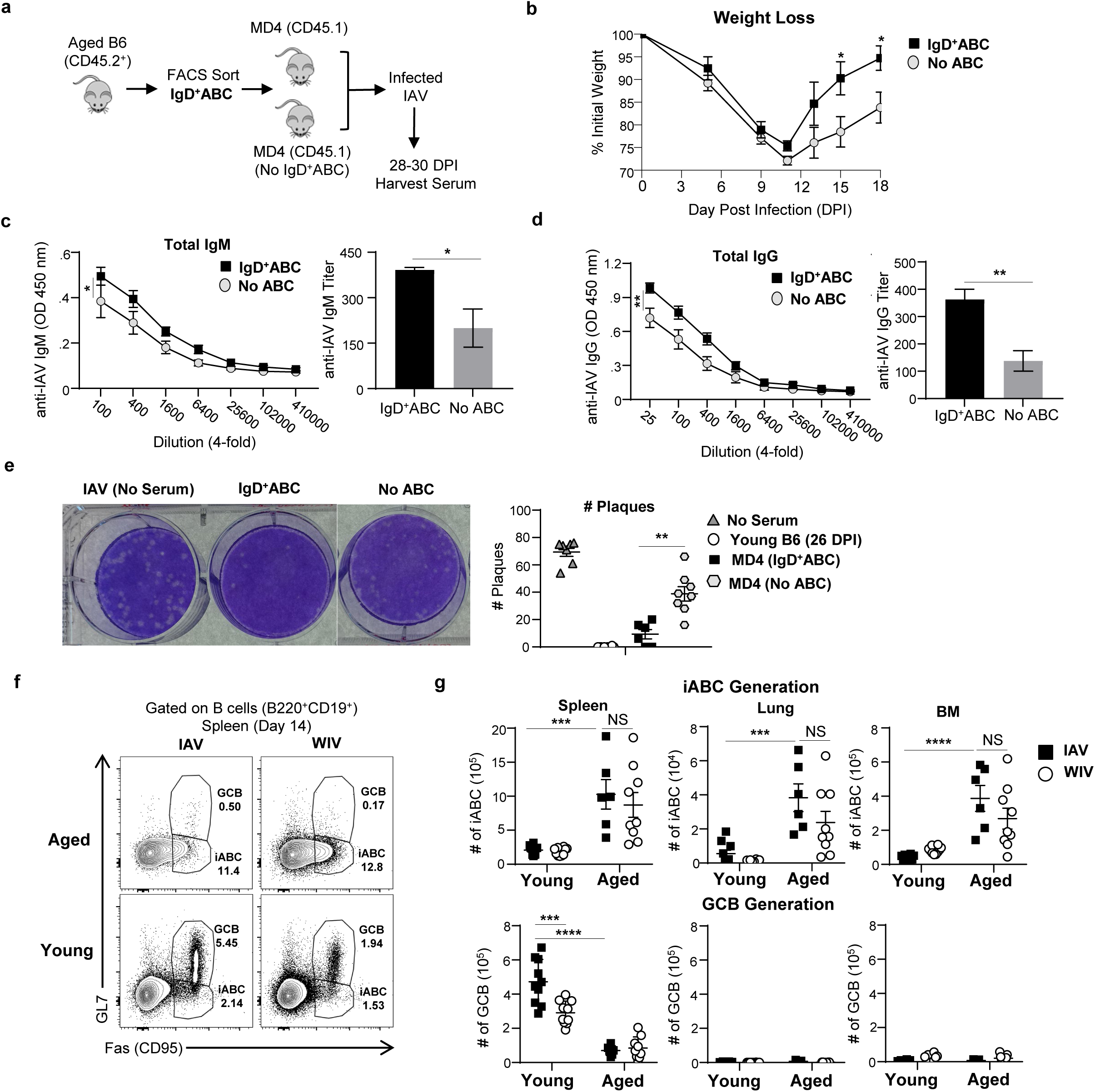
IgD^+^ABC produce neutralizing anti-IAV Ab and Inactivated IAV immunization generates iABC. (a) IgD^+^ABC (CD45.2) were sorted by flow cytometry and transferred into CD45.1 MD4 hosts and infected with 0.3 LD50 PR8 (IAV). Hosts were sacrificed and serum was collected at 28-30 dpi (b) Kinetics (0-18 dpi) of weight loss between MD4 host with IgD^+^ ABC and without IgD^+^ ABC ELISA were performed with 28-30 dpi. Results reflect serum from individual mice in both groups. (c)The titer of anti-IAV specific IgM Ab was determined by ELISA. OD curves are shown over the titrated range (left) and the titer of anti-IAV Ab (right) (d) The titer of anti-IAV specific total IgG Ab OD curves is shown over the titrated range (left) and the titer of anti-IAV Ab (right) (e) Plaque Neutralization Assay: using serum from the MD4 host with transferred IgD^+^ABC vs no ABC (n=5-6 pooled from 2-3 separate experiments) Shown is an example of the plaque assay (left) and all the data from individual serum samples (right).Induction of iABC by IAV infection vs WIV immunization. Experimental Design: Aged and young B6 mice were infected with 0.3 LD50 PR8 (IAV) i.n. or treated with 5μg whole inactivated PR8 (WIV) i.v. and sacrificed at 14 dpi. (f) Representative FACS plots showing iABC and GCB responses in the spleen of aged and young B6 mice (g) The iABC (top) and GCB response (bottom) responses in spleen, lungs and BM from individual mice in each group and is shown. Error bars represent s.e.m. Statistical significance determined by two-tailed, unpaired Student’s t-test; \**p* < .05; \*\**p* < .01; \*\*\**p* < .001, \*\*\*\**p* < .0001.

### ABC also generate an iABC response to inactivated influenza immunization

Compared to younger adults, the aged respond poorly to many current influenza vaccines due to declining CD4 and CD8 T cell responses as well as B cell responses (Sasaki et al. 2011; McElhaney et al. 2016; Nipper et al. 2018; Dugan et al. 2020). We showed that ABC can mount a T-cell independent iABC response to live IAV in aged mice (Fig .2) and that they depend on endosomal TLR signaling from host cells (Fig. 5). Whole inactivated influenza virus (WIV) contains all influenza components including many B cell epitopes, and studies indicate they can stimulate TLR7 and TLR9 (Onodera et al. 2016; Fink et al. 2018). Therefore we compared the ability of WIV immunization and live IAV infection, to stimulate iABC generation in intact aged and young mice as in our earlier studies (Brahmakshatriya et al. 2017). We examined B cell responses at 14 dpi, so we could assess both iABC and GCB development in spleen, lung and BM. In aged mice, WIV immunization or IAV infection generated equivalent numbers of iABC in the spleen, lung and BM. As expected in young mice neither immunization nor infection generated iABC, but both generated a GCB response in the spleen (Fig. 6f,g). These results suggest that WIV can provide sufficient Ag and TLR signals to drive ABC to become iABC and provide proof of principle that vaccine approaches like WIV, may succeed in harnessing ABC responses in the aged.

## Discussion

The inability to effectively vaccinate older individuals against life-threatening pulmonary infections is a major public health concern. We showed that aged mice (18-20 mo) do not mount an effective GCB response following primary influenza infection, but instead mount a novel B cell response, consisting of a prominent Fas^+^ GL7^-^ (iABC) population. These iABC are derived from a naïve subset of IgD^+^ABC, which include AbSC that produce IgM and some IgG Ab capable of neutralizing IAV. Moreover, we found that immunization with WIV as a vaccine, elicited a comparable iABC response in aged mice as live infection. Thus, it seems likely WIV provides high enough Ag and PR signals to generate iABC in the aged. Inactivated vaccines have been successfully used in the aged for decades and thus are a feasible and safe approach to induce immunity in the aged. Generation of iABC by WIV immunization is thus a first step in determining how we might harness ABC responses as an alternate source of AbSC that protect elderly populations from new pathogens or strains.

We found that the iABC generated after influenza infection expressed mostly IgM and were found in the spleen, lung and BM. This distribution resembles that seen for other T-independent responses of B1 and marginal zone B cells earlier in life, which mostly produce extrafollicular IgM^+^ Ab responses and also generate memory B cells (Cerutti et al. 2013; Alugupalli et al. 2004; Martin et al. 2001; Allman et al. 2019). Studies show that IgM^+^ memory B cells have a diverse repertoire and contribute to long-lasting durable protection (Allman et al. 2019; Bohannon et al. 2016). In young mice, primary B cell responses that migrate to the lung early following primary influenza infection are also GC-independent, predominantly IgM, and are especially protective against respiratory infection (Onodera et al. 2012). This IgM Ab can neutralize influenza virus and they are maintained over 18 months in infected and immunized mice, predicting this isotype is capable of providing durable protection (Skountzou et al. 2014; Bohannon et al. 2016). Since iABC are predominantly IgM^+^, we propose that in the aged, iABC could be a source of Ab-mediated protection following exposure to novel or emerging infections, replacing the conventional B cell response.

The iABC generated by infection in aged mice contained a small number of IgG2b^+^AbSC in the spleen, lung, and BM and some IgA^+^ AbSC cells in the lung and BM. This suggests that the iABC response drives a significant, though modest, level of isotype-switching of iABC. In young mice, influenza infection generates high levels of IgG2b and IgA, which contribute to Ab-medicated protection (Miyauchi et al. 2016; Abreu et al. 2020). Young IgG2b Ab generation has been found to be independent of TFH, but still depends on IFNγ secreted by TH1 cells (Miyauchi et al. 2016). We previously found that when we provided TLR-activated APC to aged mice, it largely restored TFH responses and led to higher total IgG titers, including increased IgG2b, in aged hosts (Brahmakshatriya et al. 2017). Given our findings here, it is likely that much of that response may have been that of ABC becoming iABC, and thus that strategies to enhance isotype-switching in iABC could potentially be developed and lead to higher IgG responses derived from ABC in the aged.

Most previous studies have not considered CD21^-^CD23^-^ABC as a heterogeneous population at different stages of development, some of which are likely to be previously Ag-experienced (Russell Knode et al. 2017; Hao et al. 2011). For instance, Cancro’s studies suggested that Ag presentation drives generation or development of CD21^-^CD23^-^ABC in unimmunized aged mice and suggests ABC are memory cells (Russell Knode et al. 2017; Cancro 2020). Moreover, most ABC studied in humans are likely memory cells and in autoimmune conditions are likely self Ag-driven (Kugler-Umana et al. 2020). However, we found that the ABC population in aged unimmunized mice, lacking most of the potential pathogen exposure, is composed of three subpopulations: a predominant IgD^+^ subset and a smaller IgD^-^ subset, consisting of IgM^+^ and IgM^-^ B cells (Fig. 2). These same three subsets were found in germ-free mice devoid of foreign and commensal Ag exposure. Thus, most IgD^+^ and IgD^-^ developed without foreign Ag exposure. Additionally, IgD^+^ABC and IgD^-^ABC respond differently to IAV infection (Fig. 3). Only the IgD^+^ population gave rise to a robust iABC in response to IAV, suggesting that they express a broad naïve BCR repertoire that includes a cohort that recognizes influenza and can respond to infection in aged mice. We conclude that IgD^+^ABC provide the largest source of protective B cells response against novel pathogens in the aged and it is thus critical to define what drives this response.

The studies in GF mice as well as their phenotype, provides strong evidence that IgD^+^ABC are a naïve subset that develops as part of an age-associated intrinsic developmental pathway (Swain et al. 2021). It is not clear whether the development of the IgD^-^ABC subsets during aging, depends on self-Ag recognition though there are studies indicating that the CD21^-^CD23^-^ABC in unimmunized SPF mice have undergone some affinity maturation compared to FOB (Russell Knode et al. 2017), suggesting that they do contain some Ag-experienced B cells. The IgD^-^ABC subset may account for those results. A strength of our study is that we have carefully traced the development of influenza Ag-experienced effectors (iABC) in aged mice from progenitor IgD^+^ABC in response to acute influenza infection.

Our studies in SAP^-/-^ hosts and T cell-depleted aged mice show that iABC generation from IgD^+^ABC does not depend on CD4 T cell help or TFH (Fig. 2, Fig. 3). The T-cell independence of the naïve ABC response, is consistent with the known loss naïve CD4 T cell response to infection and to vaccines with age (Haynes & Swain 2006). We note, that the autoimmune ABC response has been reported to require IFNγ and IL-21 produced by effector CD4 T cells (Manni et al. 2018; Naradikian, Myles, et al. 2016; Wang et al. 2018), which contrasts with the T-independent ABC expansion, indicating ABC responses in different contexts may be driven by distinct pathways. The autoimmune ABC response thus may require an immune environment dependent on CD4 T cell help, unlike anti-pathogen ABC responses, which we postulate depend on higher pathogen and Ag-presentation signals to clear of replicating virus. Further definition of strength and duration of signals needed to drive autoimmune ABC vs. protective pathogen-induced iABC response, will clearly be needed to develop safe but effective vaccines that harness the protective IgD^+^ABC in the aged.

T-cell independent B cell responses rely more on PR pathways for their generation (Bortnick & Allman 2013; Bortnick et al. 2012). Many studies have shown that cell-intrinsic TLR signaling plays an important role in somatic hypermutation and isotype class switching in conjunction with help from TFH, in GCB responses (Browne 2012). Early activation of extrafollicular autoimmune B cells in SLE, which includes different ABC subsets (Rubtsov et al. 2011; Jenks et al. 2018; Naradikian, Hao, et al. 2016), depends on cell-intrinsic TLR7 signaling (Fillatreau et al. 2020; Rawlings et al. 2017; Teichmann et al. 2013; Hua & Hou 2013). Hence, we asked whether endosomal viral-sensing TLR signals, known to be induced by influenza RNA, are required for iABC generation. We found that iABC do not develop in response to influenza infection, in either aged TLR7^-/-^ or TLR9^-/-^ mice indicating that both TLR are required for iABC generation (Fig. 5). Additionally, we found iABC development from IgD^+^ABC depends on cell-extrinsic TLR pathways, indicating non-ABC intrinsic pathways, play key role in iABC generation (Fig. 5) (Pelka et al. 2018). Our studies do not rule out a role of cell-intrinsic TLR signaling in IgD^+^ABC activation, but we predict they play a role given the previous studies (Hua & Hou 2013; Fillatreau et al. 2020). We postulate that high dependence of the ABC response on both TLR7 and TLR9, and on cell-extrinsic endosomal TLR expression, limits their response to those stimulated by pathogens that provide strong stimulation to viral sensing PR pathways (Kugler-Umana et al. 2020; Swain et al. 2021). In general, we suggest this serves as an evolutionary strategy to prevent unnecessary inflammation and the development of autoimmunity in the aged (Swain et al. 2021).

Since IgD^+^ABC become AbSC (Fig. 3), we tested whether the ABC-derived Ab can provide some level of protection against influenza (Fig. 6). We transferred 10^6^ polyclonal naïve IgD^+^ABC cells, which represents a small proportion of the aged ABC population and likely represents a small fraction of an *in-situ* ABC response in intact mice, into MD4 hosts. We found that IgD^+^ABC increase both IAV-specific IgM and IgG serum Ab titers in infected MD4 hosts and these Ab a could neutralize IAV in a plaque assay (Fig. 6). The presence of ABC in IAV-infected MD4 mice also induced faster recovery from weight loss. Thus, the increase in anti-IAV serum Ab after transfer of ABC has a modest, but significant effect on IAV clearance. Therefore, we postulate that ABC responses against influenza are a critical part of protection against novel infectious pathogens in the aged. If they become influenza specific LLPC in the BM and memory B cells in the lung, which we have not yet established, they could provide substantial local and systemic protection.

Our data suggests that ABC progenitors develop during aging as an adaptation to mount immunity against novel pathogens via iABC effector responses. In contrast to other naïve T and B immune responses whose function declines drastically with age, the ABC response seems inherently specialized to respond in an aged environment. Defining each of the factors required for protective ABC responses should lead to important insights into how to formulate and deliver vaccines to generate ABC-mediated protection and improve in anti-pathogen immunity in the highly vulnerable aged populations.

## Acknowledgements

We gratefully acknowledge the expert technical assistance of Karen Flores for maintenance of germ-free mice in the Gnotobiotic Core at the College of Veterinary Medicine, North Carolina State University. The Gnotobiotic Core is supported by the NIH funded Center for Gastrointestinal Biology and Disease, NIDDK P30 DK034987. We thank the UMMS flow cytometry core for help with cell sorting. We also would like to thank Mike Perkins for assistance with experiments and animal husbandry. Support for these studies was provided to Swain by NIH (R37AI22125, PO1AG01743, PO1AG021600, R37AG025805, PO1AI46530, R21AI1288, R21AG058758, R01AI11882 T32 AI007349, R25 GM113686 and T32 AI132152 to O.K.U.

## Data Availability Statement

Data available in article supplementary material

## Conflict of Interests

The authors declare that they have no competing interests.

## Author Contributions

O.K.U., P.D. and S.L.S. wrote the manuscript with assistance from A.M.R. and S.L.T. S.L.S. and O.K.U conceived the project. O.K.U. and P.D. designed and analyzed the experiments. O.K.U carried out experiments with help from P.D., W.Z., Y.K., J.L. C.H.C. and S.L.T. All authors contributed to editing, read and approved the submitted version.

## MATERIALS AND METHODS

### Mice

Young C57BL/6 (B6), BALB/c, and B6.CD45.1 mice were obtained from the Jackson Laboratory. Aged (8-12 weeks) C57BL/6 (B6), BALB/c, B6.CD45.1 mice were obtained from the National Institute of Aging (NIA). SAP^-/-^.CD45.1 and MD4.CD45.1 mice were bred and maintained in standard specific pathogen free (SPF) housing at the UMMS facility. BALB/c.TLR7^-/-^, BALB/c TLR9^-/-^ and Unc93b^-/-^ mice were obtained from Dr. Ann Rothstein and these strains were bred and maintained at the UMMS animal facility. Young and aged germ-free C57BL/6 mice were bred and maintained in the Gnotobiotic Core at the College of Veterinary Medicine, North Carolina State University. Germ-free mice were housed in flexible film isolators and provided with autoclaved food and water. Germ-free status was evaluated at least once a month by culturing stool samples in aerobic and anaerobic conditions. Aged SPF and GF mice were at least 72 weeks old and young mice were at least 8-wk-old prior to use. Animal use protocols were approved by the IACUCs at UMMS and at North Carolina State University.

### Virus stocks, infections and immunizations

Influenza A viruses (IAV) A/Puerto Rico/8/34 (PR8), originally from St. Jude Children’s Hospital, kindly provided by Dr. Peter Doherty, were grown and maintained at the Trudeau Institute. IIV:Formalin-inactivated influenza vaccine (A/PR/8/34 [H1N1]) was purchased from Charles River Laboratories (material no. 10100782) and used at a dose of 5 µg intravenously as in our previous studies (Xia et al. 2020). Mice were anesthetized with isoflurane (Piramal Healthcare) and were infected intranasally with influenza virus corresponding to a 0.2-0.3 LD50 dose of IAV in 50 µL of PBS.

### Adoptive B cell transfer and T cell depletion

Aged splenocytes were depleted of RBC and CD19^+^B220^+^ cells were enriched by positive selection of CD43 and Ter-119 cells and LD MACS columns (Miltenyi Biotec). CD43^-^Ter-119^-^ splenocytes were stained with FITC anti-B220 (Biolegend) BV421 anti-CD19 (Biolegend), PerCP Cy5.5 anti-CD23 (eBiosciences), APC-fluor 780 anti-CD21/35 (eBioscience), PE-Cy7 anti-IgD (Biolegend), APC anti-CD43 (eBioscience), and APC anti-CD93 (AA4.1, eBioscience). CD23^+^(FOB), IgD^+^ and IgD^-^CD21^-^CD23^-^ (ABC) lymphocytes were sorted on a BD FACS Aria III (BD Biosciences) at UMMS Flowcore. Flow cytometric sorting yielded enrichments between 90% and 95% purity for each subset. 10^6^ cells of each sorted subset were transferred intravenous into SAP^-/-^ or MD4 transgenic hosts and host were sacrificed, and spleen, BM and serum were harvested at 21-30 DPI.

In some experiments young and aged mice were treated i.p. with 250 μg of either anti–Thy1.2-depleting antibody (30-H12) (Bio X Cell) or with an isotype control on 0 DPI and 7 DPI(Strutt et al. 2012). Anti-Thy1.2 or isotype-control treated mice were infected with 0.3LD50 dose of IAV, sacrificed at 14 dpi and spleens were harvested to analyze B cell responses.

### Flow cytometry

Cells from spleen, lung and BM were harvested and passed through a 70uM nylon mesh, washed, and stained in FACS buffer [0.5% Bovine Serum Albumin, 0.01% sodium azide (Sigma-Aldrich) in PBS]. Cells were stained with amine reactive viability dyes to exclude dead cells (Invitrogen) and were blocked with anti-FcR (2.4G2) and NMS (normal mouse serum). Surface antigens were stained with fluorochrome conjugated antibodies. Antibodies used: Alexa 700 anti-CD19 (6D5), FITC B220, PerCp Cy5.5 CD23, APC-efluor 780 CD21/35, IgD(11-26c), BV650 IgM, PE CD95 (Fas, Jo2), APC GL-7, and BV421 CD138. Following surface staining, cells were fixed and permeabilized with BD Cytofix/Cytoperm (BD Bioscience) kit following manufacturer’s protocol for intracellular staining of FITC IgA (Southern Biotech), IgG2b, IgG2a/c, and PE-Cy7 IgG1isotypes (Biolegend). Antibodies were obtained from eBioscience, Biolegend, or BD Bioscience. Stained cells were acquired on an LSRII flow cytometer (BD) and analyzed using FlowJo analysis software.

### ELISA

MaxiSORP plates (Nunc) were coated overnight at 4 degrees with influenza PR8 in carbonate buffer, washed with PBS containing .05% Tween and blocked with PBS containing 1% BSA and .01% Tween for 1 hour at 37 degrees. Serum samples serially diluted in PBS-Tween PBS containing 1% BSA and .01% Tween and incubated for 2 hrs at 37^0^ C. After washing, HRP-conjugated Abs specific for mouse IgM and total IgG (Southern Biotechnology Associates) and plates were incubated 1 hr at 37^0^ C. After washing, TMB buffer (thermo) was added, and color development was stopped with 2M sulfuric acid solution. The OD reading of the color reaction was measured at 492 nm. EC50 IgM and total IgG titers were defined by half of the OD reading of the lowest dilution.

### MDCK Culture

MDCK (source) cells were seeded at 0.25×10^6^ cells(Gross et al. 2017) in a T-75flask and incubated 37 °C in 5% CO2 for 2 days or until they reach 80%-90% confluency. Cells were then washed with PBS, incubated with Trypsin at 37°C for 10 mins and FBS was added to inactivate trypsin. Cells were washed and seeded at 0.25×10^6^ cells in T-175 flasks. MDCK cells were allowed to reach 80%-90% confluency and seeded at 0.7×10^6^/well in 6 well plates for plaque neutralization assay

### Plaque Neutralization Assay

6-well plates seeded with MDCK cells were allowed to reach 120% confluency. Heat inactivated serum was serially diluted and PR8 was diluted at 150 PFU/ml in 1% Bovine Albumin Fraction V (BSA), 100 U/mL penicillin, 100 µg/mL streptomycin, 100 µg/mL CaCl/MgCl2 solution in PBS. Viral and serum dilutions were mixed and incubated at 37^0^C for an 1hr(Gross et al. 2017). 6-well plates were washed, inoculated with virus-serum mixture, and incubated at 37^0^C for 2 hr. Plates were rocked every 15 minutes to prevent drying of MDCK cells. 2 mls of agar overlay medium composed of 0.6% agarose, 0.01% DEAE Dextran, 0.099% NaHCO3, 1 µg/mL TPCK Trypsin, 50% MEM and 18% H20 solution were added to each well and the plate was incubated at 37^0^C for 48 hrs. After plaques are formed, plaques were fixed with a 4% formaldehyde solution and incubated at room temperature for 1hr. Plaques were stained with a crystal violet solution for 15 mins at room temp and plaques were counted.

## Figure Legends

**Supplemental Figure 1:**
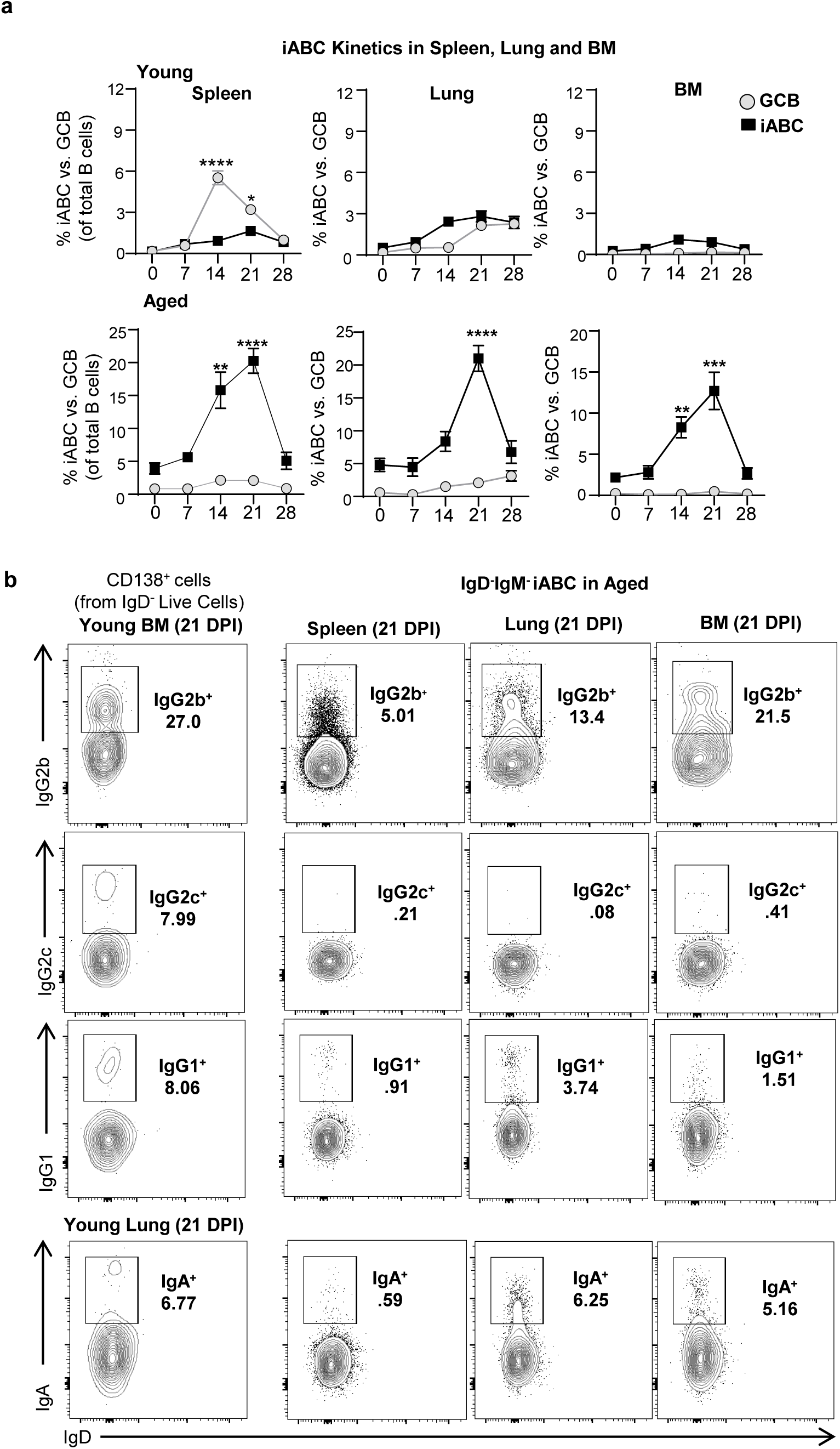
iABC and GCB response in Aged vs. Young mice. **(a)** Percent iABC (Fas^+^GL7^-^) and GCB (Fas^+^GL7^+^) gated on B cell (B220^+^CD19^+^) in young and aged mice in Spleen, Lung and BM **(b)** Representative FACS plots of IgM^+^, IgG2b^+^, IgA^+^, IgG1^+^, and IgG2c^+^ iABC in the spleen, lung and BM of infected aged mice at 21 dpi (n=6-12 pooled from 2-3 separate experiments). Gates are also shown on IgD^-^ CD138^+^ cell in BM of young mice as a positive control for gating. Error bars represent s.e.m. Statistical significance determined by two-tailed, unpaired Student’s t-test; \**p* < .05; \*\**p* < .01; \*\*\**p* < .001, \*\*\*\**p* < .0001

**Supplemental Figure 2:**
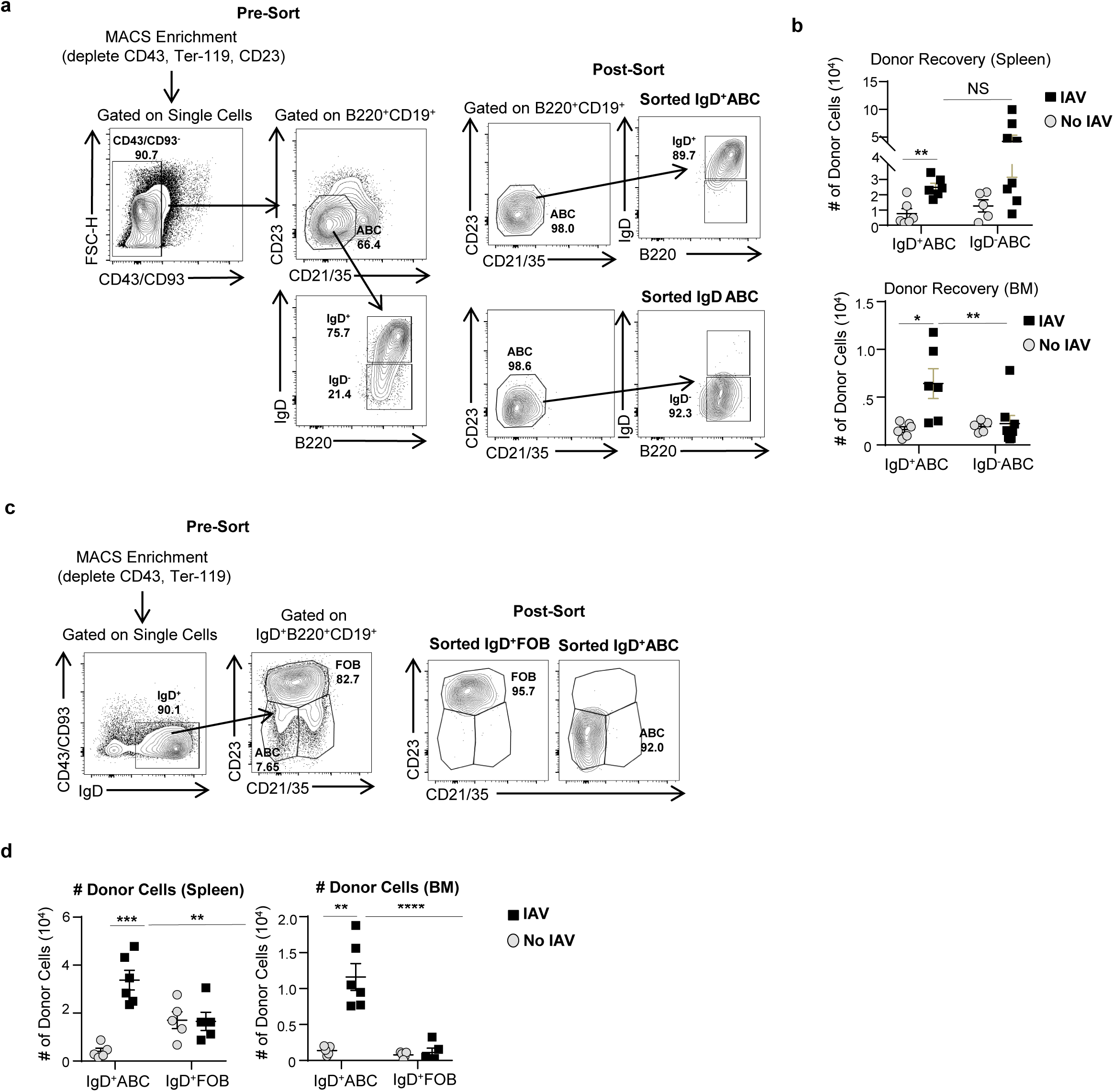
IgD^+^ABC, IgD^-^ABC, and IgD^+^FOB Purity and Donor Cell Recovery. (a) FACS Plots showing sorting strategy and purity of sorted IgD^+^ vs. IgD^-^ ABC (b) Total donor cell numbers of IgD^+^ vs. IgD^-^ABC recovered from spleen and BM of infected vs. uninfected SAP^-^/^-^ hosts (n=5-6 pooled from 3-4 separate experiments) (c) FACS Plots showing sorting strategy and purity of sorted IgD^+^ABC vs. IgD^+^FOB (d) Total donor cell numbers of IgD^+^ABC and IgD^+^ FOB, recovered from spleen of infected vs. uninfected SAP^-^/^-^ hosts. (n=5-6 pooled from 3-4 separate experiments) Error bars represent sem. Statistical significance determined by two-tailed, unpaired Student’s t-test; \**p* < .05; \*\**p* < .01; \*\*\**p* < .001, \*\*\*\**p* < .0001

**Supplemental Figure 3:**
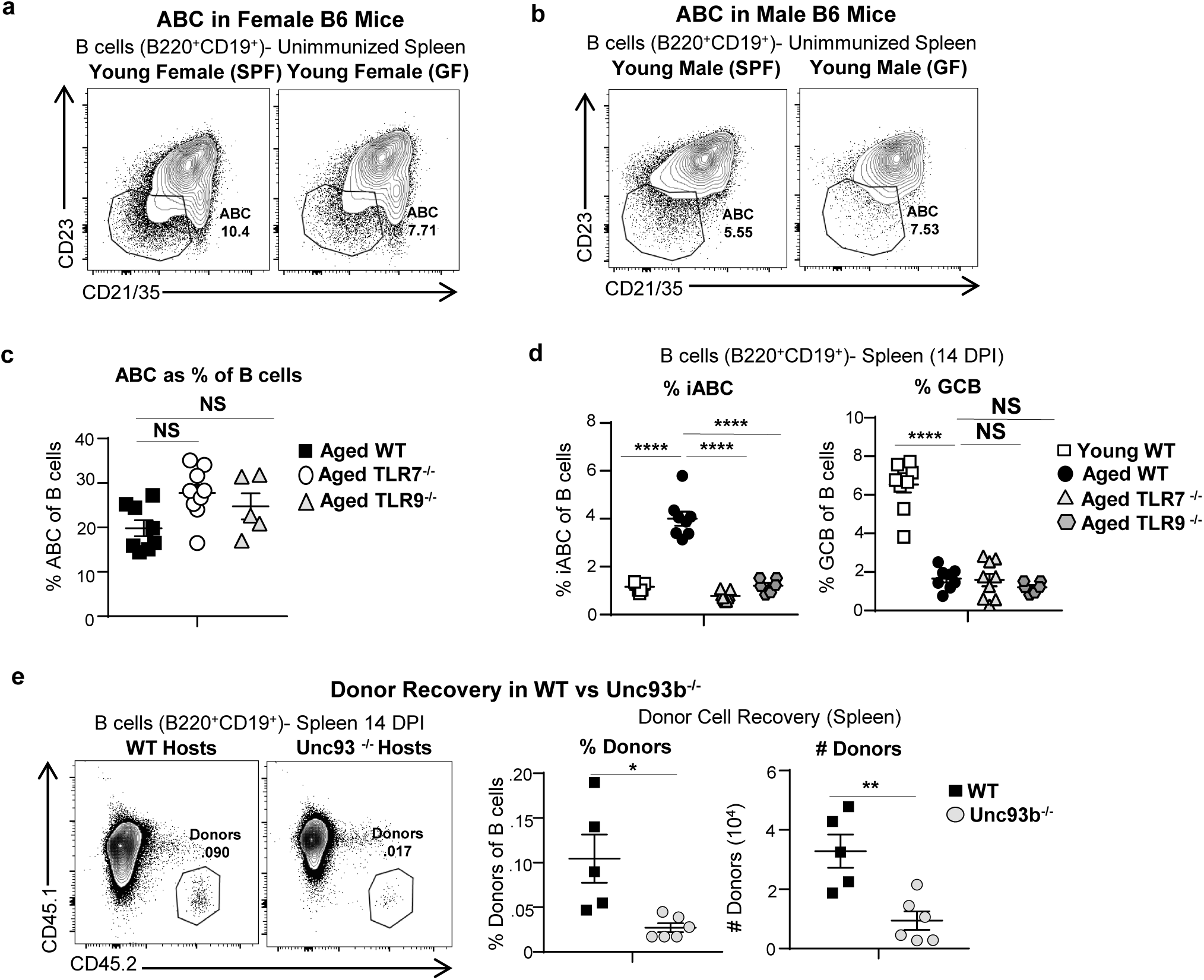
Young SPF and GF and male mice develop few ABC, TLR7 and TLR9 do not play a role in ABC Formation and Donor Cell Recovery is impaired in IAV-infected Unc93b ^-/-^vs. WT Host. (a) Representative FACS plots of total ABC (CD21^-^CD23^-^) in young SPF vs. GF B6 female mice and of (b) male mice (n=9-12 pooled from 7-8 separate experiments). (c) Graph of percent ABC from B cells in WT, TLR7^-/-^ and TLR9^-/-^ aged mice (d) Graph of percent iABC and GCB from B cells in young and aged WT, TLR7^-/-^ and TLR9^-/-^ aged mice (e) Representative FACS plots, percentage, and cell numbers of total CD45.2 IgD^+^ ABC donors in the spleen of infected CD45.1 WT and Unc93^-^/^-^ hosts. (n=5-6 pooled from 2-3 separate experiments) Error bars represent s.e.m Statistical significance determined by two-tailed, unpaired Student’s t-test; \**p* < .05; \*\**p* < .01; \*\*\**p* < .001, \*\*\*\**p* < .0001

## Notes

### Competing Interest Statement

The authors have declared no competing interest.

